# Mapping Transgene Insertion Sites Reveals Complex Interactions Between Mouse Transgenes and Neighboring Endogenous Genes

**DOI:** 10.1101/389676

**Authors:** Mallory A. Laboulaye, Xin Duan, Mu Qiao, Irene E. Whitney, Joshua R. Sanes

## Abstract

Transgenic mouse lines are routinely employed to label and manipulate distinct cell types. The transgene generally comprises cell-type specific regulatory elements linked to a cDNA encoding a reporter or other proteins. However, off-target expression seemingly unrelated to the regulatory elements in the transgene is often observed, and sometimes suspected to reflect influences related to the site of transgene integration in the genome. To test this hypothesis, we used a proximity ligation-based method, Targeted Locus Amplification (TLA), to map the insertion sites of three well-characterized transgenes that appeared to exhibit insertion site-dependent expression in retina. The nearest endogenous genes to transgenes HB9-GFP, Mito-P, and TYW3 are *Cdh6, Fat4* and *Khdrbs2,* respectively. For two lines, we demonstrate that expression reflects that of the closest endogenous gene *(Fat4* and *Cdh6),* even though the distance between transgene and endogenous gene is 550 and 680 kb, respectively. In all three lines, the transgenes decrease expression of the neighboring endogenous genes. In each case, the affected endogenous gene was expressed in at least some of the cell types that the transgenic line has been used to mark and study. These results provide insights into the effects of transgenes and endogenous genes on each other’s expression, demonstrate that mapping insertion site is valuable for interpreting results obtained with transgenic lines, and indicate that TLA is a reliable method for integration site discovery.

## INTRODUCTION

The invention of a method for generating transgenic mice by injection of a plasmid into the fertilized oocyte (Gordon and Ruddle, 1981; Brinster et al., 1981; Brinster and Palmiter, 1985) was a transformative advance in biology. In most cases, the plasmid encodes a cDNA linked to regulatory elements (promoter and enhancer) that direct its expression. These mice have been used in three main ways. In one, the purpose is to identify regulatory sequences that govern temporal and spatial patterns of gene expression; here, the cDNA encodes a reporter gene that enables expression to be mapped. In the second, the purpose is to analyze the roles of a gene product by expressing it, or a protein that interferes with it; here, the cDNA encodes the protein under study and the regulatory elements are chosen to promote the desired expression pattern. In the third, the reporter is used to mark and analyze cells that express the gene from which the regulatory elements are derived. In all three cases, the expectations are (a) that regulatory elements will direct expression in some or all of the cells in which the parent gene is normally expressed and (b) that all transgenic lines established from the same plasmid will exhibit qualitatively similar expression patterns. Indeed, these conditions are frequently met. In some cases, however, expression does not correspond to that expected from the regulatory elements included in the transgene, and/or expression patterns vary among lines. Such unexpected labeling patterns can be an advantage or a detriment. Most often, they foil attempts either to map enhancers, mark cells, or interfere with a biological process in a desired manner. They can also, however, provide unanticipated opportunities to define and mark cell types that had been undiscovered or inaccessible (e.g., Weis et al., 1991; Kim et al., 2010).

What accounts for these unpredictable expression patterns? When patterns are similar among lines established from the same plasmid, the likely explanations are that juxtapositions among normally separate regulatory elements or isolation of such sequences from their native context lead to new specificities (Swanson et al., 1985; Donoghue et al., 1991; Rao et al., 1996). In contrast, when expression patterns differ among independently generated lines, variations are generally presumed to reflect influences of endogenous sequences near the chromosomal site of integration (Palmiter et al., 1983) and are therefore termed “integration site-dependent.” The simplest explanation is that the reporter comes to be controlled by regulatory elements of a nearby endogenous gene, as seen in “enhancer traps” in transgenic Drosophila (Bier et al.,1989), zebrafish (Golling et al., 2002) and, recently, mice (Shima et al., 2016), all of which incorporate a minimal, tissue non-specific promoter but no strong regulatory elements. This mechanism is unlikely to provide a full explanation for conventional transgenes, however, which generally include tissue-specific promoters and enhancers. Other possibilities include novel specificities generated by juxtapositions of transgene and endogenous sequences, variations in chromatin conformation near the integration site, differences in transgene copy number, or mutations of either the transgene or flanking sequences that occur upon transgene integration (Palmiter and Brinster, 1986; Feng et al., 2000).

Having generated many transgenic lines with such insertion site-dependent expression patterns (Weis et al., 1991; Feng et al., 2000; Kim et al., 2010), we have become interested in the relationship between insertion sites and transgene expression patterns. In a few cases, the expression of a transgene has been related to that of a specific, nearby endogenous gene (Kothary et al., 1988, Sharpe et. al, 1999; Narboux-Nême et al., 2012). We reasoned that if this were a general phenomenon, the endogenous gene might play a role in the development or function of the marked cells. Unfortunately, although identifying insertion site is straightforward for invertebrates, available methods for mice (Burgess et al., 1995; Sharpe et. al, 1999; Suzuki et al., 2006; Sha et al., 2007; Liang et al., 2008; Dubose et al, 2013; Srivastava et al., 2014; Raman et al., 2015) have been cumbersome, little used and, in our hands, largely unsuccessful. Recently, however, a newly developed method termed Targeted Locus Amplification (TLA) was introduced that appeared to be more promising (de Vree et al., 2014; Cain-Hom et al., 2017). In TLA, genomic DNA in nuclei is cross-linked by formaldehyde, digested into small fragments by the frequently cutting NlaIII restriction enzyme and religated to form larger circular DNA containing fragments that were likely to have been near neighbors on a chromosome. These fragments are de-crosslinked and digested by another restriction enzyme, NspI, to create ~2kb fragments, which are then subjected to PCR with primers derived from sequences unique to the transgene. By amplifying fragments that contain the transgene sequence, this step selectively amplifies sequences from neighboring fragments. The product is sequenced and mapped to the genome, thereby localizing the transgene and also revealing insertions, deletions or other structural rearrangements both within the transgene and in flanking sequences.

We used TLA to determine insertion sites for three transgenic lines that incorporate fluorescent proteins as reporters: HB9-GFP (green; Wichterle et. al, 2002), Mito-P (cyan, CFP; Misgeld et al., 2007), and TYW3 (yellow, YFP; Kim et. al, 2010). All label subsets of cells in retina by what appears to be an insertion site-dependent mechanism, and have been used in studies of retinal development and function (Schubert et. al, 2008; Kim et al., 2010; Trenholm et al., 2011; Kay et al., 2011a; Kay et al., 2011b; Kay et al., 2012; Duan et al., 2014; Krishnaswamy et al., 2015; Shekhar et al., 2016; Peng et al., 2017; Sethuramanujam et al., 2017; Ray et al., 2018). For two of them, our interest was heightened by breeding experiments detailed in Results, which suggested that the transgenes were linked to genes expressed in some of the retinal cells marked by the fluorescent protein: *Cdh6* in one line and *Fat4* in another. For all three, we document interactions between the transgene and the closest endogenous gene. For two of them, the nearest endogenous gene is hundreds of kilobases (kb) from the transgene and yet it appears to strongly influence transgene expression. For all three, the transgene decreases expression of a nearby endogenous gene in a position-dependent manner. Together, our results provide novel insights into insertion site-dependent transgene expression and strengthen the argument that determination of insertion sites can be useful both for gene discovery and for assessing effects of transgene insertion that would otherwise go undetected.

## METHODS

### Animals

Animal protocols were approved by the Institutional Animal Care and Use Committee (IACUC) at Harvard University. Animals were used in accordance with NIH guidelines. Mutants were maintained on a C57/B6J background. We obtained both the HB9-GFP (Wichterle et al, 2011) and the Thy1-mitoCFP-P (Misgeld et al., 2007) transgenic mouse lines from Jackson Laboratories. For brevity, we refer to Thy1-mitoCFP-P as Mito-P. The *Fat4* conditional mutant (Saburi et. al, 2008) was a kind gift of Helen McNeill (U. Toronto). The TYW3 line was generated in our laboratory using a Thy1-lox-YFP-STOP-lox-WGA-ires-LacZ sequence, as previously reported (Kim et al., 2010). The Cdh6^CreER^ line was also generated in house by targeted insertion of a frt-neo-frt cassette, a 6xMyc-tagged CreER-T2, and a poly-adenylation signal at the translational start site of the *Cdh6* coding sequence (Kay et. al, 2011a).

### Histology

Mice were euthanized by intraperitoneal injection of Euthasol. Eyes were removed and fixed in 4% PFA in PBS for 90 minutes. Retinas were then dissected and rinsed with PBS. Retinas to be sectioned were sunk in 30% sucrose in PBS overnight at 4°C, embedded in tissue freezing medium, frozen in dry ice and stored at -80°C until processing. Retinas were then sectioned at 20μm on a cryostat. Sections were rehydrated in PBS, incubated in 5% Normal Donkey Serum (NDS), 0.3% Triton X-100 in PBS for 2 hours and then incubated with primary antibodies overnight at 4°C. Sections were then washed in PBS, incubated with secondary antibodies for 2 hours at room temperature, washed again, dried, and mounted with Vectashield (Vector Lab).

For whole mounts, fixed retinas were incubated with 5% NDS, 0.1% Triton X-100 in PBS for 3 hours and then incubated in primary antibody for 5 days at 4°C. Retinas were then washed in PBS and incubated overnight in secondary antibody. Finally, retinas were washed in PBS, flat-mounted on cellulose membrane filters (Millipore), coverslipped with Fluoro-Gel (Electron Microscopy Sciences), and sealed.

Antibodies used were as follows: rabbit and chicken anti-GFP (1:2000, Millipore; 1:1000, Abcam); goat anti-choline acetyltransferase (ChAT) (1:500, Millipore); mouse anti-TFAP2 (1:200, DSHB); guinea pig and rabbit anti-Rbpms (1:2000, PhosphoSolutions; 1:300, Abcam). Rabbit and guinea pig antibodies against Slm1 (1:5000) and Slm2 (1:2500) were the generous gift of Peter Scheiffele (Iijima et al., 2014). Secondary antibodies were conjugated to Alexa Fluor 488 (Invitrogen), Alexa Fluor 568 (Invitrogen), or Alexa Fluor 647 (Jackson ImmunoResearch) and used at 1:1000. Nuclei were stained with ToPro Cy5 (1:5000, Thermo Fisher).

*In situ* hybridization was performed as described elsewhere (Kay et al., 2011; Duan et al., 2014). Tissue was collected and prepared with RNase-free reagents, sectioned and imaged as described above. Section hybridization was carried out at 65°C. Probes were detected using anti-digoxigenin (DIG) antibodies conjugated to horseradish peroxidase (HRP), followed by amplification with Cy3-tyramide (TSA-Plus System; Perkin-Elmer Life Sciences, MA) for 2hrs.

Images were acquired using 488, 568, and 647 nm lasers on an Olympus-FV1000 Confocal Microscope. We used ImageJ (NIH) software to analyze confocal stacks and generate maximum intensity projections.

### Targeted Locus Amplification

Targeted Locus Amplification (TLA) technology uses the physical proximity of nucleotides within a locus of interest to generate a map of original sequences and corresponding inserted transgenes (de Vree et al., 2014). Transgenic homozygotes and wild-type controls were euthanized and cells were prepared from their spleens (Cain Hom et al., 2017). Homozygotes were distinguished from heterozygotes by fluorescent quantitative PCR (qPCR) results from a commercial genotyping service (Transnetyx; http://www.transnetyx.com/). In some cases we confirmed their results in our laboratory by mating or additional PCR. The cells were frozen and shipped to Cergentis (Utrecht, the Netherlands). TLA was then performed as described in de Vree et al. (2014) and Hottentot et al. (2016). Briefly, DNA was crosslinked, fragmented, religated, and decrosslinked. This product served as the TLA template, which was subsequently fragmented, circularized, and amplified with inverse primers complementary to a short locus-specific sequence. Once the complete locus was amplified, ~2kb segments were sheared. Libraries were prepared for sequencing by MiSeq or HiSeq technologies on an Illumina platform.

### PCR of Genomic DNA

Genomic PCR was used to confirm insertion sites and deletions revealed by TLA. Primer sequences were as follows:

Primer 1 : 5’ - AACTTGTGCGGTTCTGTCCT -3’
Primer 2: 5’- TT GACAAAGT GGGGGTTAGGC-3’
Primer 3: 5’- CAGCGAAGGGGAAATTTGCATAT-3’
Primer 4: 5’ -GCAT CT GT GT GT CACAGC AGT GGT -3’
Primer 5: 5’ - CTAGCCAAAGGGATT AACAAT GT G-3’
Primer 6: 5’ - CAAT CAT AAT GCA GA CA GGAAT GT -3’
Primer 7: 5’ - CA GA GCT CT GGGT CCA GT CA GT A-3’
Primer 8: 5’- TATGTGCGCCACTGTGTAGTT-3’
Primer 9: 5’- TTTGGTTCCCGGTCTCTGAAG -3’
Primer 10: 5’ - AT CCT GTT GCAGCGT CGTTA-3’
Primer 11: 5’ - GT CAGGGACCT CT GT GGTT G-3’

We confirmed the predicted insertion of HB9-GFP on Chromosome 15 using two primer pairs, one for the left junction Primers 1 and 2, and one for the right junction, Primers 3 and 4. We confirmed the predicted insertion of Mito-P on Chromosome 3 with Primers 5 and 6 and the predicted deletion in Chromosome 3 with Primers 5 and 7. We confirmed the insertion of TYW3 with Primers 8 and 9 and the predicted deletion in Chromosome 1 with Primers 10 and 11.

We extracted DNA from tail-clips of wild-type, heterozygous, and homozygous animals with 50μL Quick Extract (Lucigen) at 68°C for 30 min and 98°C for 3min in a PCR machine. Zygosity was determined as described above. PCR reaction mixtures were 3μL DNA, 12.5μL Econotaq Plus Green 2X Master Mix, 7.5μL H_2_0, 1μL F primer 10mM, 1μL R primer 10mM for 25μL reactions. The reaction program was 94°C for 2 min; [94°C for 30s, 50-55°C for 30s, 72°C for 1 min] x 40 cycles; 72°C for 5 min. The annealing temperature varied in accordance with the melting temperature of the primer pairs tested.

### RT qPCR

Mice were euthanized by intraperitoneal injection of Euthasol. Retinas were dissected and RNA was extracted with 250μL Trizol Reagent. Reverse transcription was performed using SuperScript III (Invitrogen). qPCR was performed using KAPA SYBR FAST qPCR master mix (Kapa Biosystems). We normalized expression of our genes of interest to Gapdh levels and ran samples in triplicate on an ABI 7900. Resulting CT values were used to calculate ΔΔCT and fold changes in the expression of endogenous genes in our three transgenic lines. Both technical and experimental replicates were included. In most case, wild-type littermates were used as controls. Primers were as follows:

*GAPDH-F:* 5’ - GT GGAGT CAT ACT GGAACAT GT AG-3’
*GAPDH-R:* 3’ -AATGGTGAAGGTCGGTGTG-5’
*CDH6-F:* 5’ - CAA GA GGCT GGA CA GGGAAG-3’
*CDH6-R:* 3’ -CGGGA CT GT GGCT GT GT AAA-5’
*FAT4-F:* 5’-GGT GCCAACGCT CT GGT CACGT AT GC-3’
*FAT4-R:* 3’-CAGGGGTT GT GT CTT CT GGGAT GT C-5’
*SLM1-F:* 5’-GCTACGTGACCCCAACACAA-3’
*SLM1-F:* 3’-CTGTCGTAGGCATCCTCGTT-5’

### Estimating Copy Number

We estimated transgene copy number from quantitative fluorescent PCR data provided by our genotyping service (Transnetyx). The raw signal returned for every sample was a function of the ΔCT between the housekeeping gene *c-Jun* and a probe for our gene of interest:

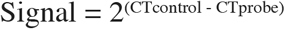

We compiled raw signal data for heterozygous animals genotyped with GFP or LacZ probes, averaged the signal for each transgenic strain. We normalized these values to the average intensity returned for our single copy Cre-GFP and LacZ knock-in lines and calculated an estimated copy number for our three transgenic lines of interest.

### Statistical Analysis

Comparisons were performed using GraphPad Prism software. For single comparisons, we used Student’s t test. For multiple comparisons, we used one-way ANOVA.

We calculated the odds of transgene and endogenous gene expression overlapping in individual cell types using elementary combinatorics with the null hypothesis being that expression is independent. Thus, given 3/120 Cdh6+ types and 2/120 GFP+ types in the retina, the odds of finding at least 1 double+ type is given as,

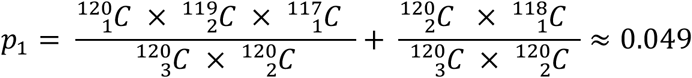

Similarly, given 2/120 Fat4+ types and 3/120 CFP+ types in the retina, the odds of finding 2 double+ type is given as,

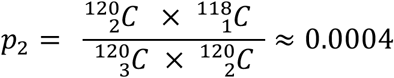

## RESULTS

### The HB9-GFP transgene is inserted near the Cdh6 locus on mouse Chromosome 15

The HB9-GFP transgene is composed of a 9 kb fragment from the 5’ end of the *Mnx1* gene (previously called Hb9) that extends into the first exon, linked to a cDNA encoding the enhanced green fluorescent protein (GFP) (Wichterle et al, 2002) (Fig. 3a). It was generated to label motor neurons, which express *Mnx1,* but was later shown to also mark two types of retinal neurons: a subset of cone photoreceptors and a type of retinal ganglion cell (RGC) that responds selectively to dark or bright objects moving in a dorsal-to-ventral direction across the retina (ventral-preferring on-off direction-selective retinal ganglion cells or V-ooDSGCs; Trenholm et al., 2011, 2013; Fig. 1a, b, e). RNA-Seq data generated in our laboratory show that neither cones nor V-ooDSGCs express *Mnx1* at detectable levels (Peng et al., 2017; Sarin et al., 2018).

**Figure 1.**
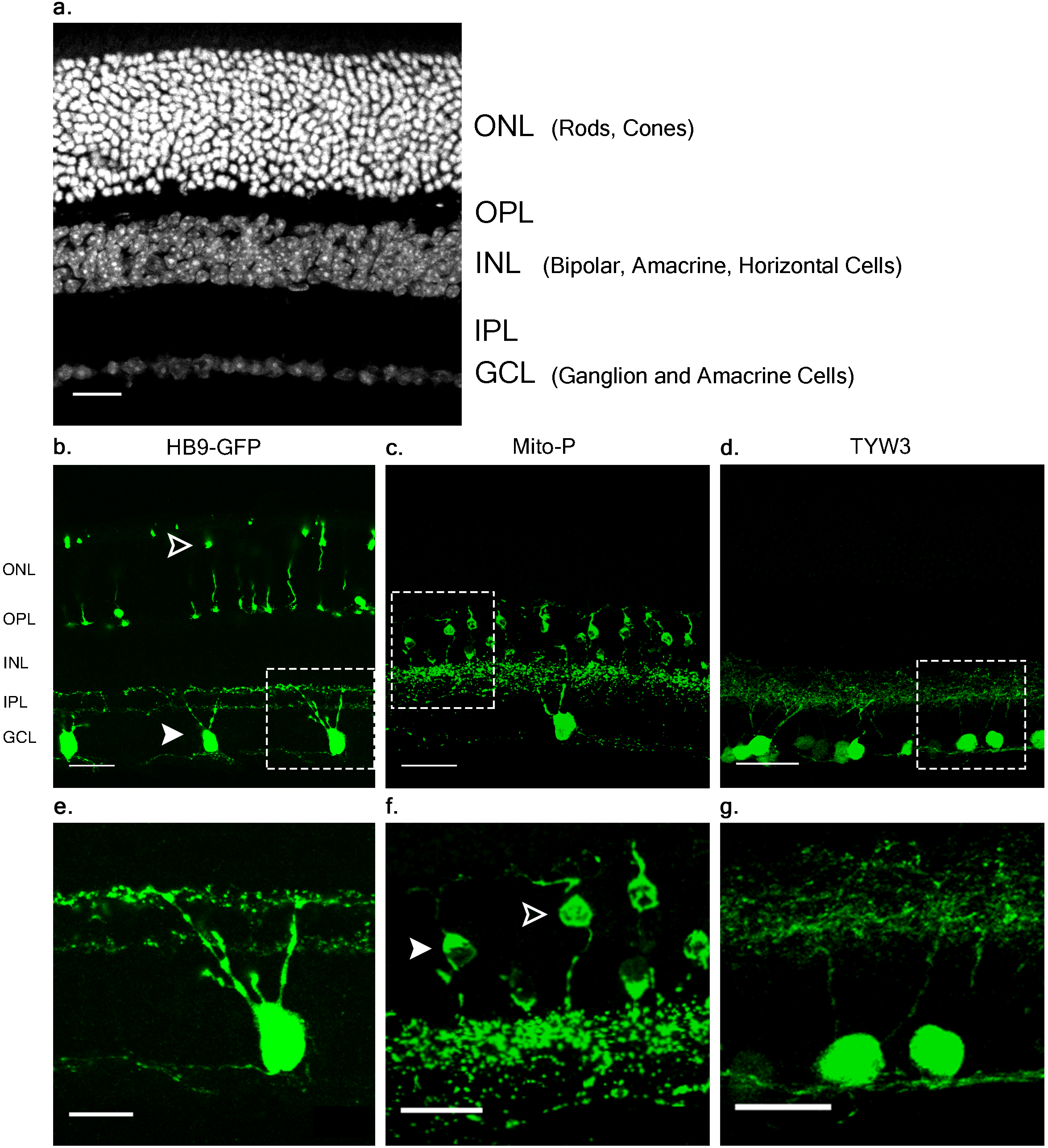
Retinal expression pattern of three transgenic mouse lines. (a) Section of an adult mouse retina stained with ToPro, a nuclear stain. Photoreceptors are located in the outermost layer, termed Outer Nuclear Layer (ONL). These form synapses in the Outer Plexiform Layer (OPL) with interneurons, whose cell bodies reside in the Inner Nuclear Layer (INL). Bipolar and amacrine cells of the INL also form synapses in the Inner Plexiform Layer (IPL), with ganglion cells from the Ganglion Cell Layer (GCL). (b, e) Expression of HB9-GFP in V-ooDSGCs (solid arrowheads) and cone photoreceptors (open arrowheads). (c, f) Expression of Mito-CFP in Type 1a (arrowheads) and 1b Bipolar Cells (BCs), as well as other bipolar, amacrine, and ganglion cells. (d, g) Expression of TYW3 in several types of retinal ganglion cells (RGCs) that stratify in the middle part of the IPL. All cells are GFP positive. Scale bars a-d are 40μm and scale bars e-g are 20μm. Boxed regions in b-d are shown at higher magnification in e-g.

In a study of ooDSGCs, we discovered that V-ooDSGCs and ventral preferring ooDSGCs (D-ooDSGCs) both express *Cdh6,* which encodes the recognition molecule Cadherin 6 (Kay et al., 2011a; Fig. 2a). *Cdh6* is also expressed in a set of amacrine interneurons, termed starburst amacrine cells, which innervate ooDSGCs but do not express HB9-GFP; conversely, cones are HB9-GFP-positive but Cdh6-negative. Thus, of >120 retinal neuronal types (M.A.L, I.E.W, J.R.S., in preparation), one expresses *Cdh6* but not HB9-GFP, one expresses HB9-GFP but not *Cdh6,* one expresses both and >115 express neither.

**Figure 2.**
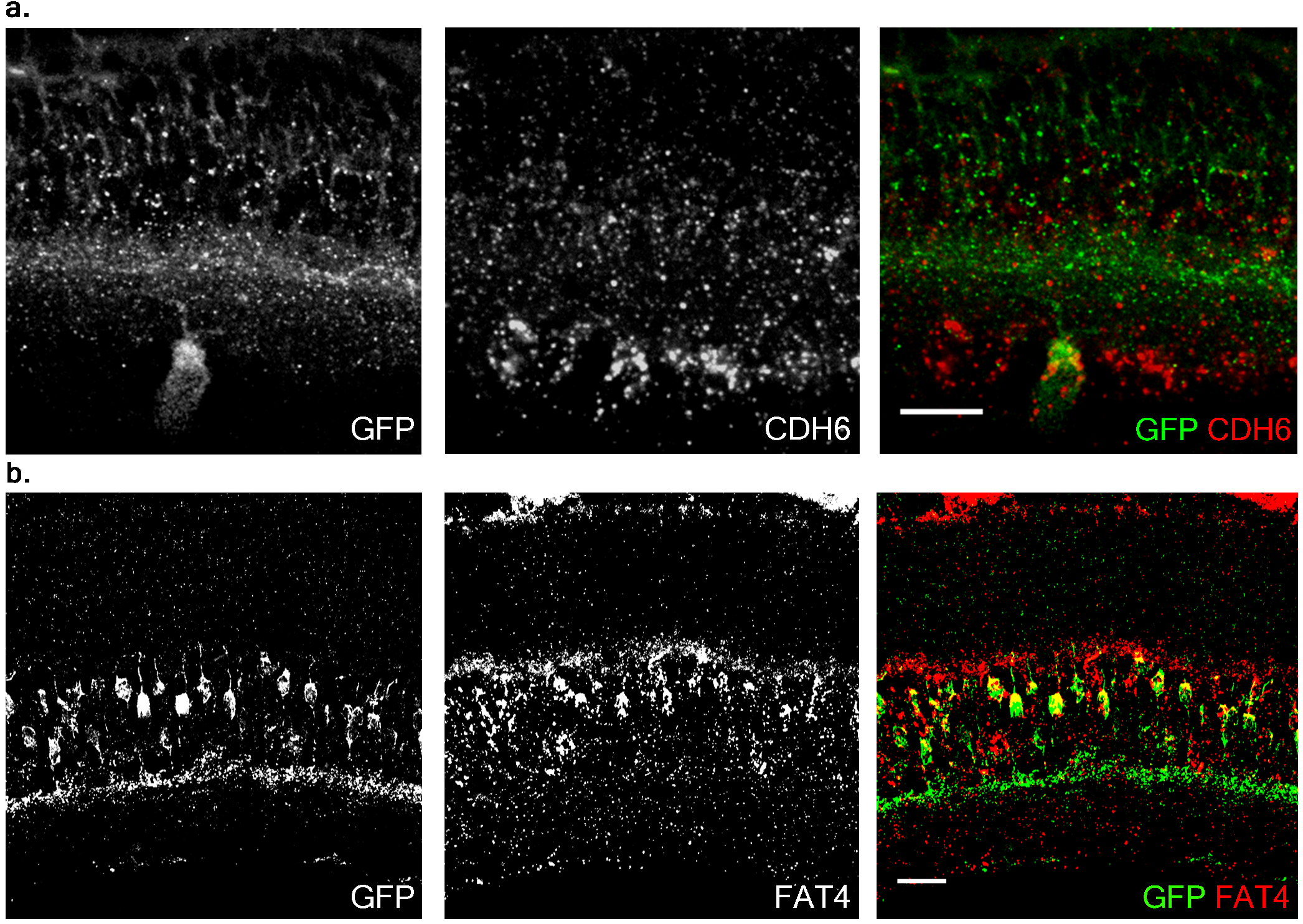
Co-expression of transgenes and endogenous genes. a) Retinal expression pattern of *Cdh6* mRNA shown by in situ hybridization. *Cdh6* is expressed in HB9-GFP positive V-ooDSGCs. (b) Retinal expression pattern of *Fat4* mRNA shown by in situ hybridization. *Fat4* is expressed in Mito-P positive BCs. Scale bars are 20μm.

To study the role of Cdh6 in the development and function of V-ooDSCGs, we generated *Cdh6* mutants and attempted to generate HB9-GFP^+/−^; Cdh6^−/−^ mice by crossing HB9-GFP^+/−^;Cdh6^+/−^ and *Cdh6^−/−^* mice. However, we retrieved no HB9-GFP^+/−^; Cdh6^−/−^ offspring, suggesting that the HB9-GFP transgene and the endogenous *Cdh6* gene were linked. We used TLA to test this possibility. Two primer pairs complementary to the transgene sequence were designed, one complementary to GFP sequences at the 3’ end of the transgene and the other complementary to *Mnx1* sequences at the 5’ end of the transgene (Fig. 3a). Both were used to generate products that were sequenced to a depth of 5Mb.

**Figure 3.**
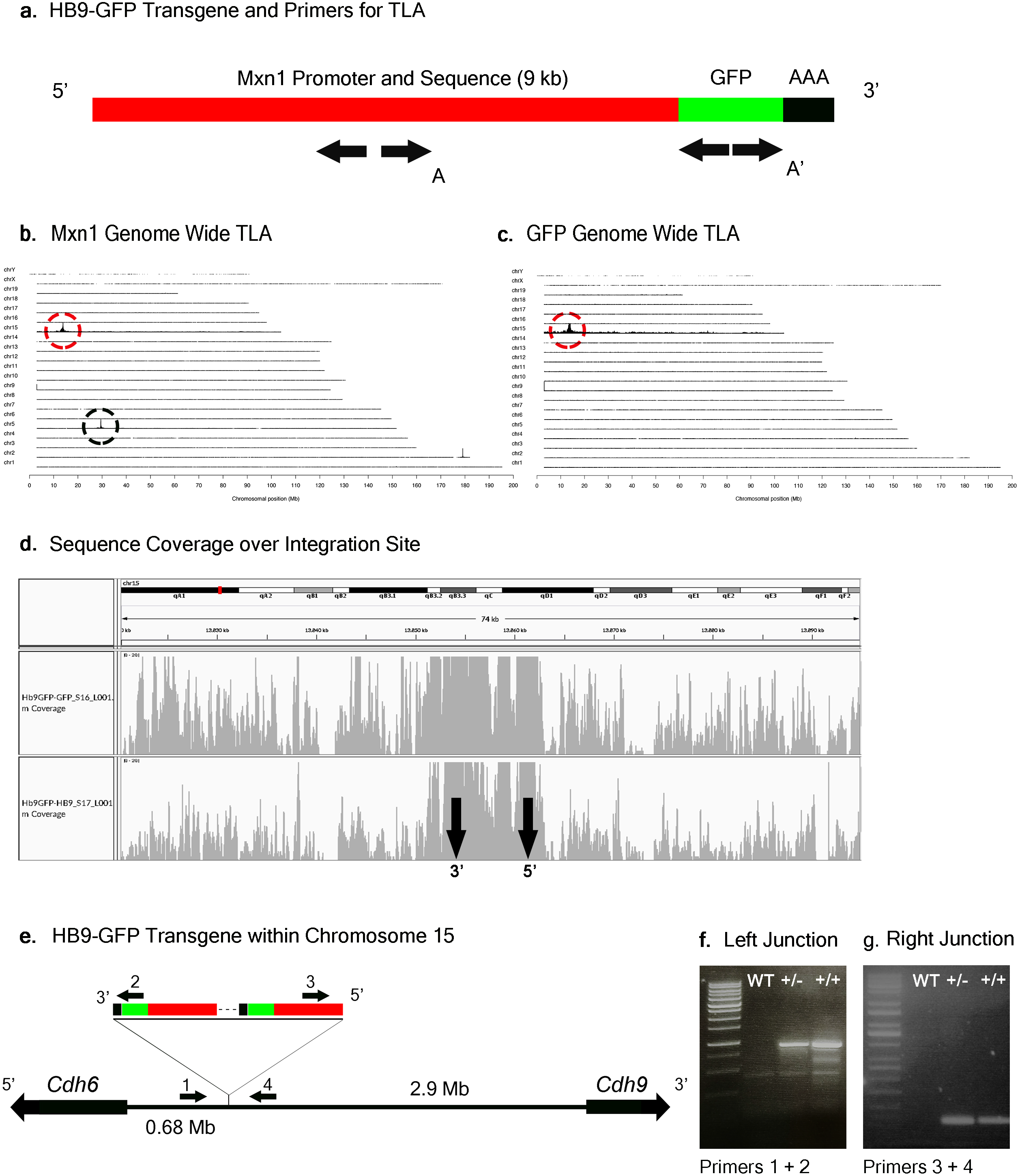
HB9-GFP Locus Identification. a) Schematic of the HB9-GFP transgene, showing positions of primers used for TLA. Primer sets were designed within the Mxn1 and the GFP sequences. (b) Genome-wide TLA coverage using Mnx1 primers. Peak at Chromosome 5 shows endogenous Mnx1 (denoted by black circle) and peak at Chromosome 15 shows inserted sequence (red circle). (c) Genome-wide TLA coverage using GFP primers, showing a peak at Chromosome 15 (red circle). (d) Regional coverage of the HB9-GFP insertion site on Chromosome 15, spanning 74 kb surrounding the integration site. (e) Schematic of the inserted sequence. Multiple copies of the transgene were inserted 3’ to 5’. HB9-GFP is located between *Cdh6* and *Cdh9* on Chromosome 15. Primers were designed to confirm the left and right junctions of the transgene with Chromosome 15. (f) Confirmed insertion of the transgene with primers specific to the junction between the 3’ end of the transgene and Chromosome 15 (Primers 1 and 2). (g) Confirmed insertion of the transgene with primers specific to the 5’ end of the transgene and Chromosome 15 (Primers 3 and 4).

Results from both sets of primers identified the insertion site of the HB9-GFP transgene on Chromosome 15 (Chr15: 13,853,116 -13,862,097) (Fig. 3b-d). Consistent with our prediction, the gene nearest to the HB9-GFP transgene was *Cdh6* (Chr15:13,034,200-13,173,675), with the 3’ end of *Cdh6* ~680 kb upstream of the 5’end of HB9-GFP. To confirm the insertion, we designed primers flanking the predicted left and right junctions. Genomic PCR confirmed the insertion of HB9-GFP in heterozygotes and homozygotes for both of these primer sets (Fig. 3f-g). Based on the orientation of the junctions and the relative position of the transgene, we conclude that the transgene was inserted 3’ to 5’. We also identified a duplication of a ~9 kb segment on Chromosome 15 within this insertion site.

### The Mito-P transgene is inserted near *Fat4* on Chromosome 3

In the Mito-P transgene, the coding sequence of the enhanced cyan fluorescent protein (CFP) was fused to a sequence encoding a 31 amino acid fragment from the human subunit VIII of the cytochrome c oxidase gene sufficient to drive expression in mitochondria (Misgeld et al., 2007). This construct was inserted into a 6.5 kb fragment of the *Thy1* gene that is known to drive expression in central projection neurons including motor and sensory neurons and RGCs (Caroni, 1997, Feng et al., 2000) (Fig. 4a). This transgene was designed and used to monitor mitochondrial dynamics in motor axons. Several lines were generated, each of which labeled distinct neuronal types, indicating an insertion site-dependent pattern (Misgeld et. al, 2007). In addition to motor axons, Mito-P also labels two of 15 types of bipolar interneurons (Type 1A and Type 1B) as well as one of ~50 types of amacrine interneuron (nGnG) in retina (Schubert et al., 2008; Kay et. al, 2011b; Shekhar et al., 2016). Bipolar and amacrine cells express *Thy1* at low levels; a small number of RGCs, which express *Thy1* at far higher levels, are also labeled (Fig. 1a, c, f).

**Figure 4.**
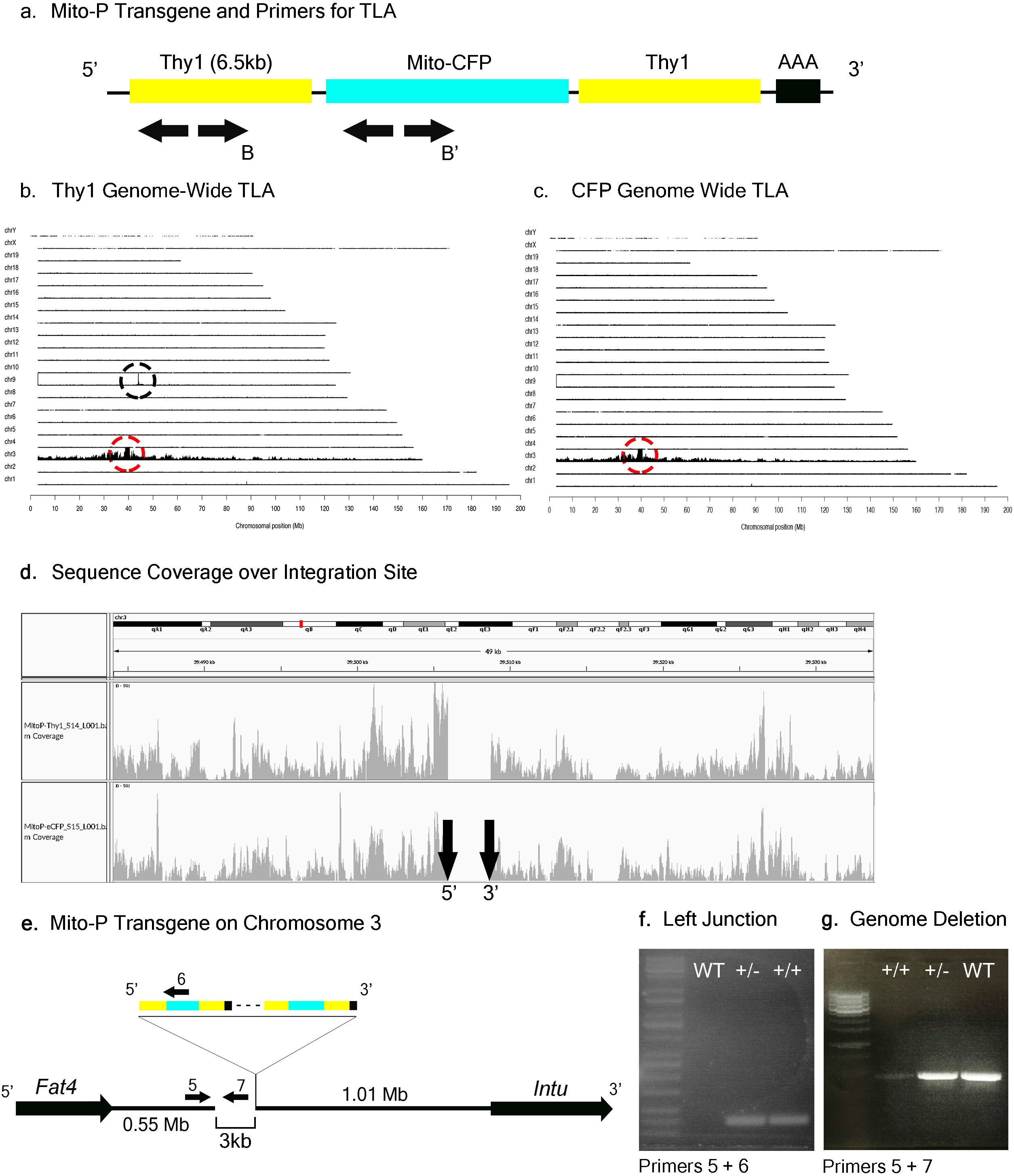
Mito-P Locus Identification. a) Schematic of the Mito-P transgene, showing positions of primers used for TLA. Primer sets were designed within the *Thy1* sequence and within the CFP sequence. (b) Genome-wide TLA coverage using *Thy1* primers. Peak at Chromosome 9 shows endogenous *Thy1* (black circle) and peak at Chromosome 3 shows inserted sequence (red circle). (c) Genome-wide TLA coverage using CFP primers, with a peak at Chromosome 3 showing the inserted sequence (red circle). (d) Regional coverage of the Mito-P insertion site on Chromosome 3 using both sets of primers, showing a 3 kb deletion and spanning 49 kb. (e) Schematic of the inserted sequence. The transgene was inserted multiple times between *Fat4* and *Intu* in Chromosome 3, in the 5’ to 3’ direction. Primers were designed to confirm the left junction of the transgene to Chromosome 3, as well as the predicted deletion. (f) Primers 5 and 6 were used to confirm the junction between the transgene and Chromosome 3 in heterozygous and homozygous animals. (g) Primers 5 and 7 were used to confirm the deletion engendered by the transgene. The band is absent in putative homozygous Mito-P animals.

In a study of mouse bipolar cells, we found that Types 1A and 1B expressed *Fat4,* which encodes a cell surface protein implicated in cell polarity (Shekhar, et al., 2016; Fig. 2b). To study the role of *Fat4* in retinal development, we obtained conditional *Fat4* mutants (Saburi et al., 2008) and attempted to generate Mito-P; *Fat4*^loxP/loxP^ mice by breeding. However, we retrieved no Mito-P;Fat4^loxP/loxP^ offspring from Mito-P;*Fat4*^loxP/+^ x *Fat4*^loxP/loxP^ matings. This result, which paralleled that described above for HB9-GFP and *Cdh6,* suggested that the Mito-P transgene was inserted near the *Fat4* locus.

TLA revealed that the Mito-P transgene was inserted in Chromosome 3 (Chr3: 39,505,947 – 39,508,740), and that the insertion was accompanied by a 3 kb deletion (Fig. 4a-d). The insertion site is located 550 kb from the 3’ end of the *Fat4* gene (Chr3: 38,886,940-38,952,429), accounting for our failure to recover Mito-P;Fat4^loxP/loxP^ offspring via conventional recombination. We confirmed both the insertion site and the accompanying deletion by PCR on genomic DNA (Fig 4e-g).

### The TYW3 transgene is inserted near *Khdrbs2* on Chromosome 1

We generated a set of transgenic mice called TYW using the *Thy1* sequences described above (Caroni, 1997; Feng et al., 2000). The transgene included a cDNA encoding YFP flanked by LoxP sites followed by cDNAs for E.Coli beta galactosidase (LacZ) and wheat germ agglutinin (WGA) (Fig. 5a). It was designed to express YFP constitutively and LacZ plus WGA following excision of the floxed cassette with Cre recombinase (Kim et al., 2010). In practice, however, LacZ and WGA were expressed at undetectable levels, but the YFP was expressed strongly. Each of several lines labeled distinct sets of retinal neurons. TYW3 labeled six of ~45 RGC types. One type was labeled most brightly; we called it W3B and analyzed its development and function in detail (Zhang et al., 2012; Krishnaswamy et al., 2015). Remarkably, dendrites of all 6 (W3B and five dimmer types together called W3D) laminated in a narrow central stratum in the central third of the inner plexiform layer (I.E.W. and J.R.S., in preparation). Assuming that all thirds of the IPL are populated by equal numbers of dendrites, the odds of 6 stratifying in the same third are (1/3^6^)x3 or 1/243. We therefore wondered whether sequences near the TYW3 insertion site contributed to this unusual expression pattern.

**Figure 5.**
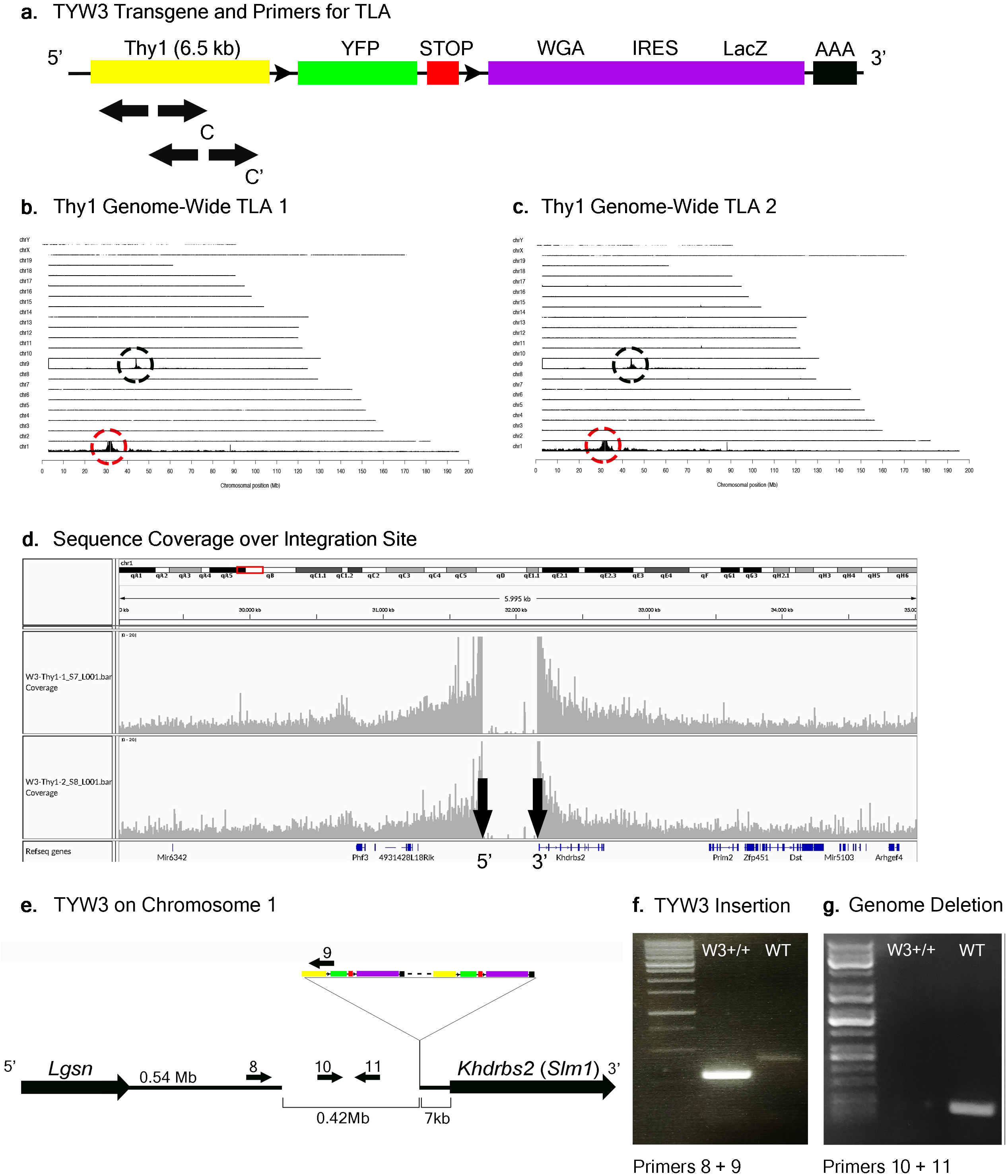
TYW3 Locus Identification. (a) Schematic of the TYW3 transgene, showing positions of primers used for TLA. Both primer sets were designed within the *Thy1* sequence. (b) Genome-wide TLA coverage using *Thy1* primer set 1. Peak at Chromosome 9 shows endogenous *Thy1* (black circle) and peak at Chromosome 1 shows inserted sequence (red circle). (c) Genome-wide TLA coverage using *Thy1* primer set 2. Peak at Chromosome 9 shows endogenous *Thy1* (black circle) and peak at Chromosome 1 shows inserted sequence (red circle). (d) Regional coverage of the TYW3 insertion site on Chromosome 1 using both sets of primers, showing a 419 kb deletion and spanning 5 Mb. (e) Schematic of the inserted sequence on Chromosome 1. The transgene was inserted multiple times between *Lgsn* and *Khdrbs2* in Chromosome 1, in the 5’ to 3’ orientation. Primers were designed to confirm the left junction between the transgene and Chromosome 1, as well as the predicted deletion. (f) Primers 8 and 9 were used to confirm the junction between the 5’ end of the transgene and the 3’ end of Chromosome 1 in homozygous animals. This 292 bp band is absent in wild-type animals (g) Primers 10 and 11 were used to confirm the deletion engendered by the transgene. The 173 bp band is absent in putative homozygous TYW3 animals.

TLA revealed that the TYW3 transgene was inserted in Chromosome 1 (Chr1: 31,745,752-32,165,062). The insertion was accompanied by a 420 kb deletion directly upstream of the insertion site (Fig. 5a-d). The 3’ end of the TYW3 transgene is 7 kb upstream of the initiation codon of the *Khdrbs2* gene (Fig. 5d, e), which encodes an RNA binding regulator of alternative splicing also called Slm1 (Ehrmann et al., 2016). We validated the insertion site with primers spanning the junction between the 3’ end of Chromosome 1 and the 5’ end of the transgene (Fig. 5f), as well as the deletion caused by the insertion using primers for a sequence within this putatively deleted sequence (Fig. 5g).

### Insertion site maps enable determination of zygosity by conventional PCR genotyping

When transgenic mice are inbred, heterozygotes are generally distinguished from homozygotes either by qPCR of genomic DNA or by outcrossing to wild type animals. The former, which we used to assess zygosity for TLA, is subject to considerable variation and the latter is cumbersome. Moreover, the use of primers derived from the reporter (e.g., GFP) can also give ambiguous results if, for example, more than one line contains a fluorescent protein as occurs in some complex mating schemes. Once insertion sites are mapped, however, line-specific primers can be designed that allow one to distinguish zygosity without relying on relative fluorescent RT-PCR intensities (Cain-Hom et al., 2017). We demonstrate this for two of the three lines analyzed here. For Mito-P, Primers 5 and 7 in Fig. 4e generate a band in wild types or heterozygotes but none in homozygotes, because the sequence recognized by Primer 7 is deleted (Fig. 4g). Likewise, Primers 10 and 11 in Fig. 5e recognize a sequence deleted in the TWY 3 line, so PCR using these primers generates a band in wild types or heterozygotes but not in homozygotes (Fig. 5g). In addition, one primer set in each case generates a band unique to the lines: Primers 1 and 2 or 3 and 4 for Hb9-GFP (Fig. 3e-g), Primers 5 and 6 for Mito-P (Fig. 4e,f) and Primers 8 and 9 for TYW3 (Fig. 5e, f). Thus, information gained from the insertion site greatly simplifies genotyping.

### All three transgenes are present in multiple copies

Multiple copies of transgenes are frequently integrated in a head-to-tail tandem array at a single genomic site (Palmiter and Brinster, 1986). Transgene copy number can vary from one to over one hundred, and the number can influence both qualitative and quantitative aspects of transgene expression (e.g., Williams et al., 2008). To more completely characterize the HB9-GFP, Mito-P and TYW3 transgenes, we estimated their copy number. To this end, we analyzed data from the commercial service (Transnetyx; www.transnetyx.com) that we employ to genotype our transgenic lines. Transnetyx uses a quantitative fluorescent PCR method to detect transgene-specific sequences. The raw quantitative PCR results are normalized to a control endogenous gene *(cJun)*. We compared the relative intensities of our three transgenic lines to intensities of single copy knock-in lines. After averaging the signals of 11-56 heterozygous animals from each line and normalizing to single copy intensities of GFP (n=20) or LacZ (n=13), we estimate that the HB9-GFP, Mito-P and TYW3 transgenes contain 10-11 (10.57±0.60), 13 (12.92±0.53) and 8 copies (8.05± 0.79) respectively (Fig. 6) (Mean ± SEM).

**Figure 6.**
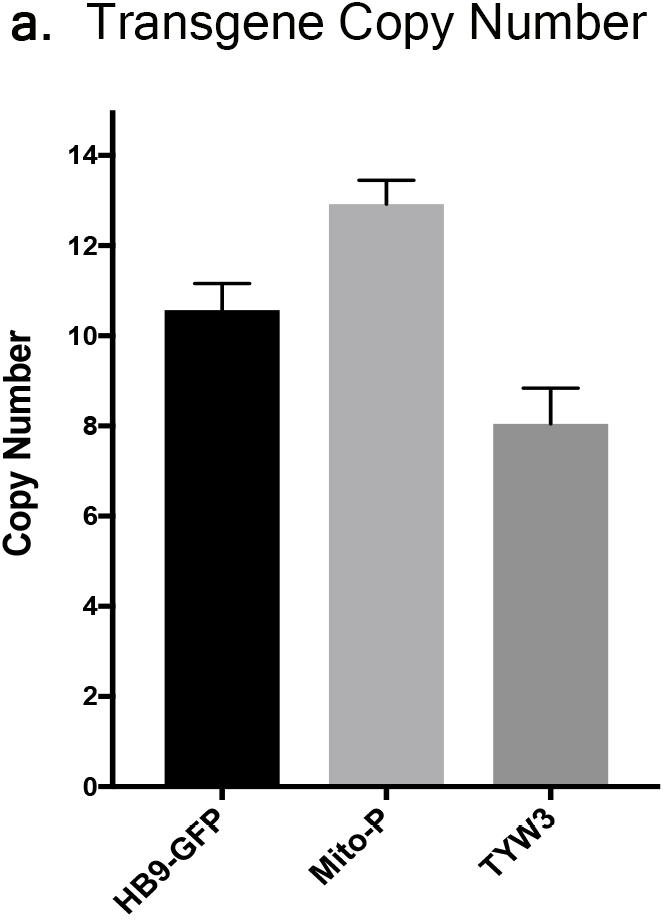
Estimation of Transgene Copy Number. Copy number was determined by quantitative PCR of genomic DNA, with values normalized to a line with a single copy of GFP or LacZ. Signal intensities were garnered from the following number of animals: n= 13-20 for single GFP and LacZ lines; n=46 for HB9-GFP; n=56 for Mito-P; n=11 for TYW3.

In each case, we are confident that all copies are inserted at a single genomic site for two reasons. First, TLA revealed only a single insertion site for each transgene (Figures 3–5). Second, even if more than a single insertion occurred initially, the lines have been bred for at least 10 years, or >40 generations, which is more than enough to segregate multiple inserts.

### Effect of transgenes on expression of neighboring endogenous genes

We next asked if the transgenes we had studied affected expression of neighboring endogenous genes. Using quantitative PCR (qPCR), we found that levels of *Khdrbs2* mRNA were reduced by ~45% in TYW3 homozygotes compared to controls; *Fat4* mRNA levels were reduced by ~25% in Mito-P homozygotes compared to controls; and *Cdh6* mRNA levels were reduced by ~10% in HB9-GFP homozygotes compared to controls. The reductions were statistically significant for all three lines (p<0.0001 for TYW3 and Mito-P; p=0.0094 for HB9-GFP by Student’s t-test). Interestingly, the effect size of these transgenes on endogenous gene expression is related to the distance between the two (Fig. 7).

**Figure 7.**
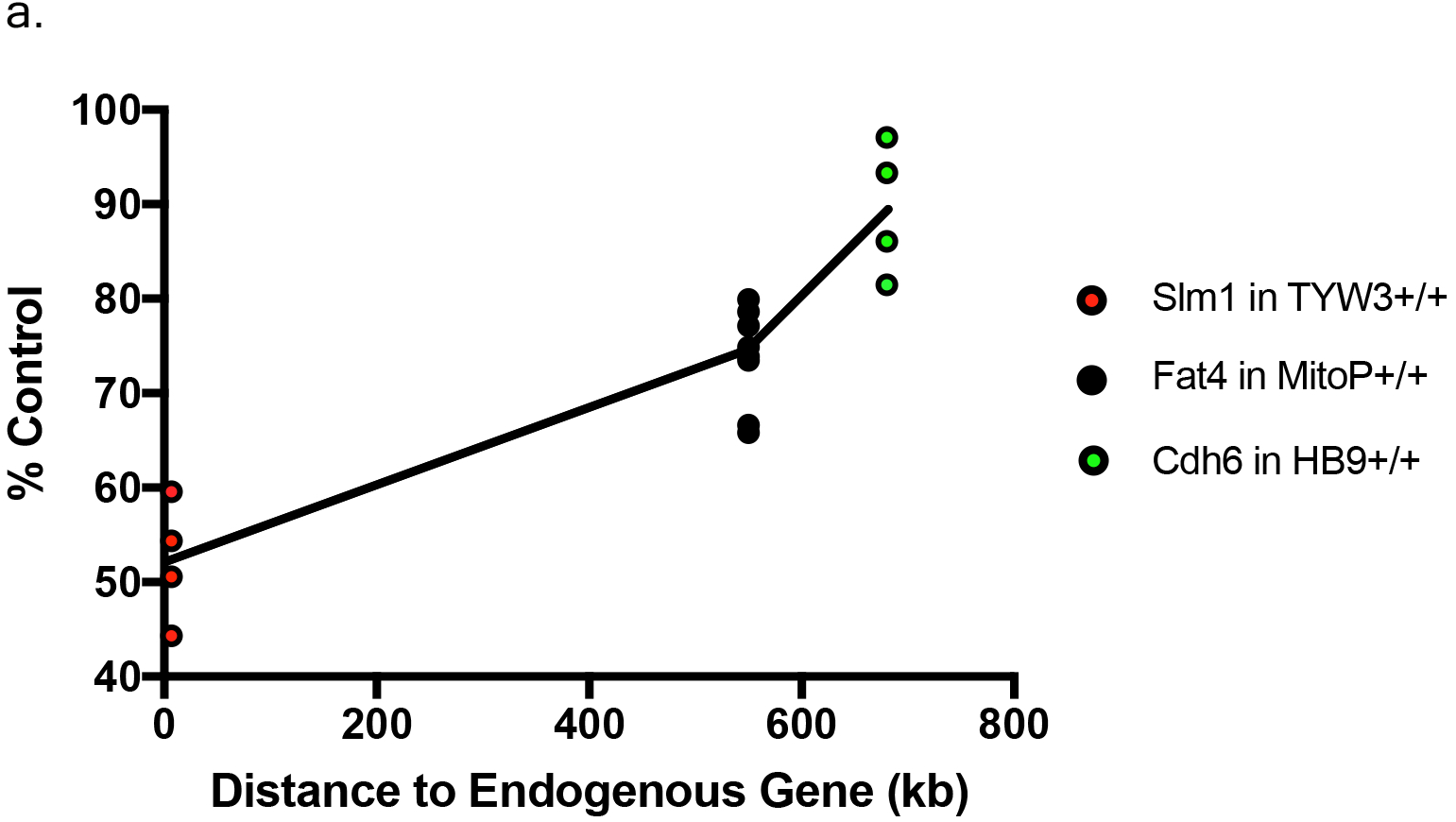
Effect of transgenes on expression of neighboring endogenous genes. Expression of *Slm1, Fat4,* and *Cdh6* mRNA from TYW3, Mito-P, and HB9-GFP homozygous animals, respectively, was determined by qPCR of RNA from adult retina. Values were compared to those from wild-type littermates for Mito-P, and HB9-GFP and to age-matched controls for TYW3. ΔΔ CT values were calculated against Δ CT values of *Gapdh.* The difference was calculated as 2^^-ΔΔct^. The change in expression compared to control was significant for all three transgenic lines: *Slm1* in TYW3 homozygotes (p<0.0001), *Fat4* in Mito-P homozygotes (p<0.0001), and *Cdh6* in HB9-GFP homozygotes (p<0.009) (HB9-GFP n=4; Mito-P n=8; TYW3 n=4). Significance was calculated by one-way ANOVA and Tukey’s multiple comparison tests. Effect of transgene on endogenous gene expression varies with distance between the transgene and the endogenous gene

### Interactions between the TYW3 transgene and the endogenous *Khdrbs2* gene

Because the TYW3 transgene exerted a strong effect on expression of *Khdrbs2,* we used immunohistochemical methods to examine interactions between the transgene and the endogenous gene in cellular detail. In wild-type retinas, Slm1 was present in most RGCs, as identified by double-labeling with the pan-RGC marker Rbpms (Rodriguez et al., 2014) and most amacrine cells, identified with the pan-amacrine marker AP2 (Bassett et al., 2007) (Fig. 8a and 8b). Slm1 also appeared to be expressed by horizontal cells, identified by soma position. Bipolar, photoreceptors, and Müller glial cells were not detectably labeled. Similar patterns of Slm1 labeling were observed in TYW3 heterozygotes, and nearly all YFP-positive RGCs, which comprise ~15% of all RGCs, were Slm1-positive (Fig. 8c).

**Figure 8.**
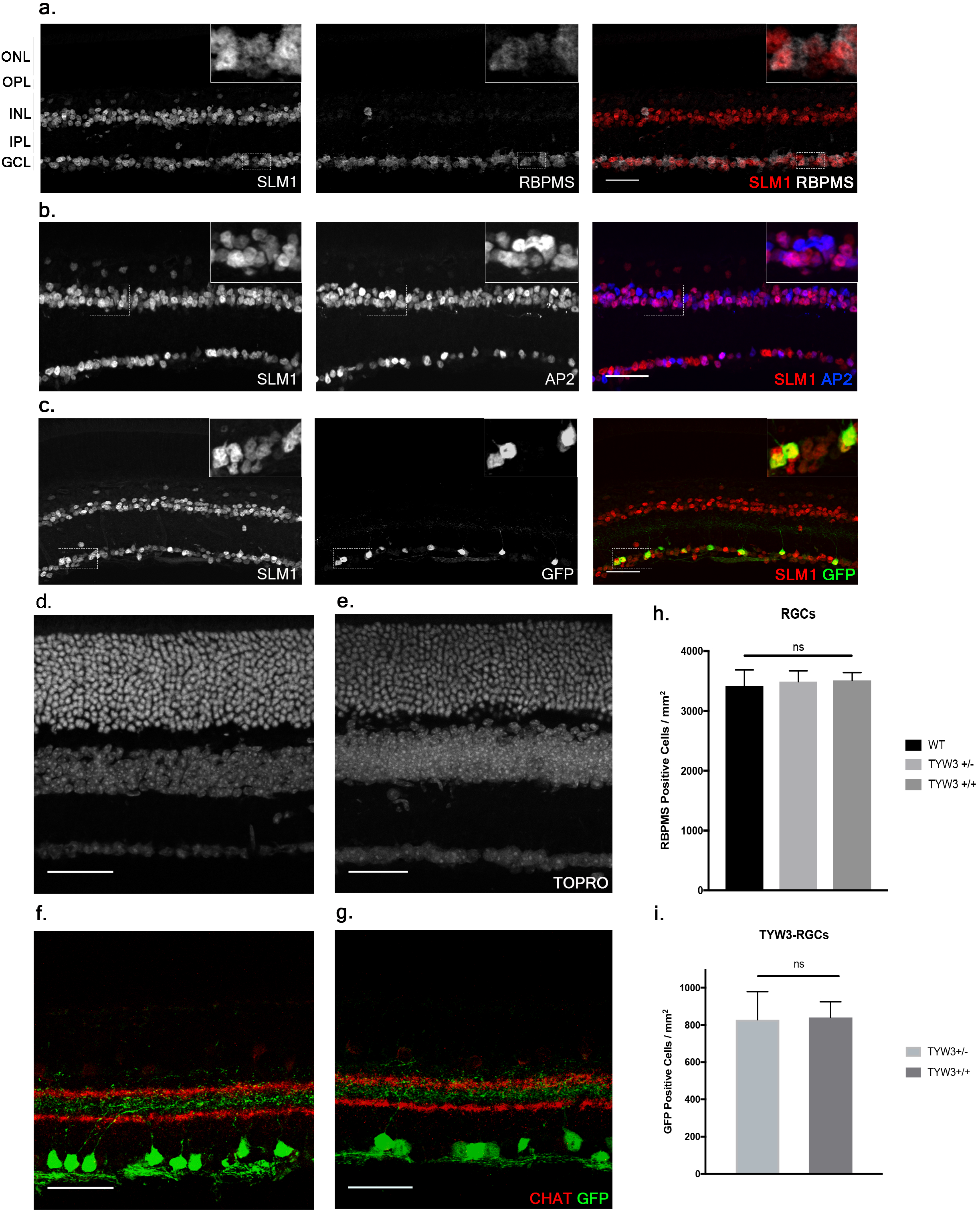
Expression of Slm1 in wild-type retina and TYW3 retinal architecture. (a) Slm1 is expressed by most RGCs. Stained with Rbpms and Slm1. (b) Slm1 is expressed by most amacrine cells. Stained with TFAP2 (AP2) and Slm1. (c) Slm1 is expressed by YFP-positive RGCs in the TYW3 line. Stained with GFP and Slm1. Sections from wild type (d) and TYW3 homozygous (e) retina. Stained with ToPro. Lamination of TYW3-RGCs in heterozygous (f) and homozygous (g) retina. Stained with GFP and ChAT, labeling starburst amacrine cells. (h) Rbpms counts in wild-type, TYW3+/-, and TYW3+/+ retinas (Mean ± SEM). No significant difference between conditions (n=5 for WT, n=4 for TYW3+/- and TYW3+/+). (i) TYW3-RGC counts in TYW3+/- and TYW3+/+ retinas (Mean ± SD). No significant difference between conditions (n=4 for both). Scale bars for a-g are 40μm.

We then assessed TYW3 homozygotes, which, as noted above, are *Khdrbs2* hypomorphs. We detected no alterations in the overall structure of the retina (Fig. 8d and e) or in the lamination pattern of TYW3 RGCs (Fig. 8f and g). We found no significant change in the total number of RGCs (Rbpms-positive) in TYW3 homozygotes (Fig. 8h). Likewise, the number of YFP-positive RGCs did not differ significantly between TYW3 heterozygotes and homozygotes (Fig. 8i).

Although there were no changes in the general organization of TYW3+/+ retinas, decreased levels of Slm1 were apparent in both RGCs and amacrine cells. We found that only 39.9 ± 3.4% of Rbpms-positive RGCs (Fig. 9a-c) and 46.8 ± 1.0 % of AP2-positive amacrines were Slm1-positive in TYW3 homozygotes (Mean ± SEM) (Fig. 9d-f). Interestingly, however, the loss of Slm1 from transgene-positive RGCs in TYW3 homozygotes was greater than that in RGCs generally: only 5.2 ± 2.3 % of W3-RGCs were detectably Slm1-positive in homozygotes (Fig. 9g-i).

**Figure 9.**
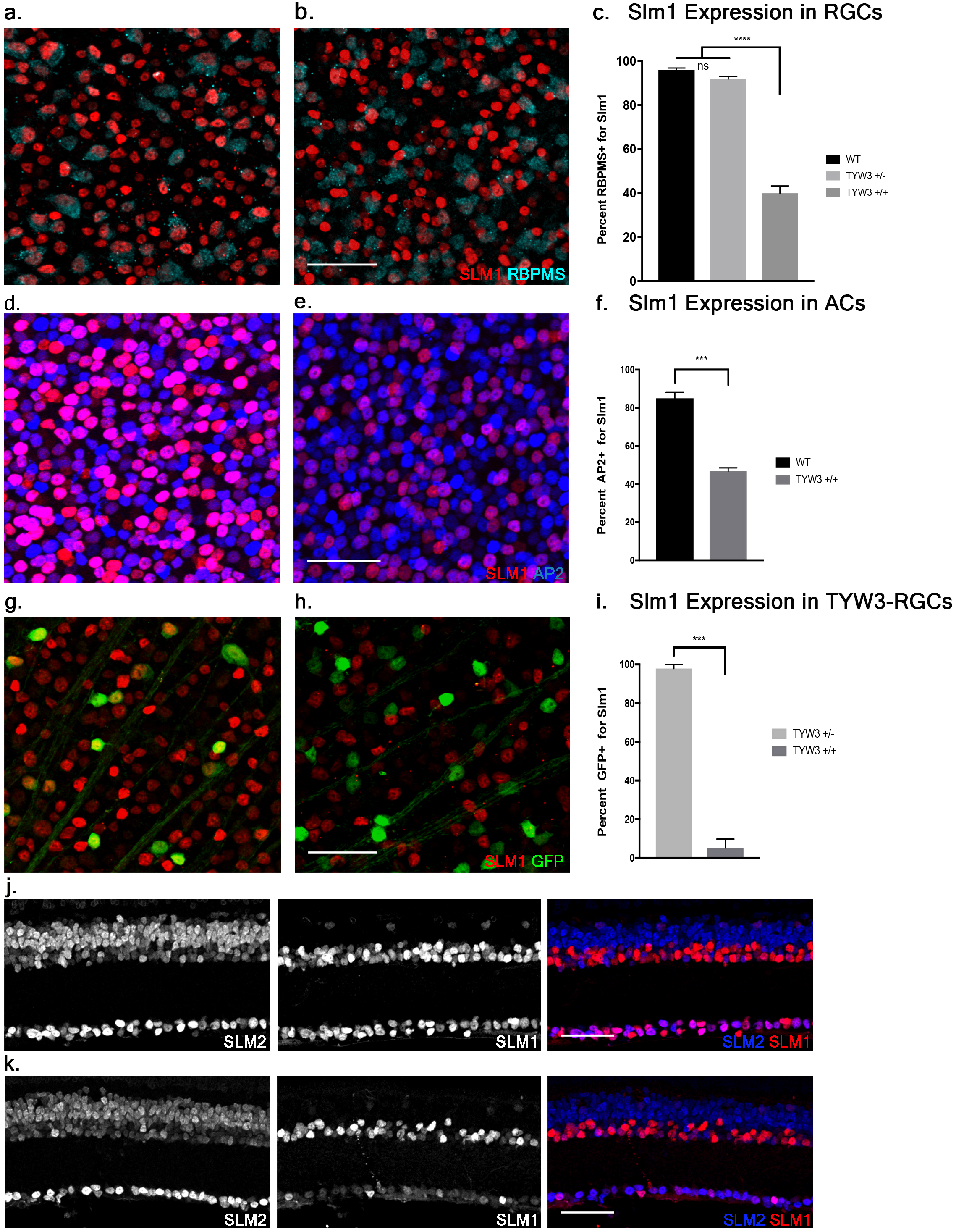
Expression of Slm1 and Slm2 in TYW3 homozygotes. Expression of Slm1 in RGCs in wholemount of wild-type (a) and TYW3 homozygous (b) retinas. Stained with Rbpms and Slm1. (c) Quantification of Slm1 expression in RGCs across TYW3 genotypes. In wild-type animals, 96.1 ± 0.8% of RGCs express Slm1, while in TYW3 homozygotes, only 39.9 ± 2.3% of Rbpms-positive RGCs express Slm1 (Mean ± SEM). There is no significant decrease in Slm1 expression by RGCs in TYW3 heterozygotes. Significance determined by one-way ANOVA (n=5 for WT, n=4 for TYW3+/- and TYW3+/+). Expression of Slm1 in amacrine cells in wholemount in wild-type (d) and TYW3 homozygous (e) retina. Stained with AP2 and Slm1. (f) Quantification of Slm1 expression in amacrine cells across TYW3 genotypes. In wild-type animals, 85.0 ± 0.9% of RGCs express Slm1, while in TYW3 homozygotes, only 46.8 ± 0.5% of AP2-positive ACs express Slm1. Significance determined by t-test (n=3 for both) (Mean ± SEM). Slm1 expression in TYW3-RGCs in wholemount in TYW3 heterozygous (g) and TYW3 homozygous (h) retina. Stained with GFP and Slm1. (i) Quantification of Slm1 expression in TYW3-RGCs. As in section, 97.9 ± 1.0 % of heterozygous TYW3-RGCs express Slm1, while in TYW3 homozygotes, only 5.2 ± 2.3 % still express Slm1. Significance determined by t-test (n=4 for both) (Mean ± SEM). (j) Slm1 and Slm2 expression overlaps in wild-type retina. Slm2 is also expressed in bipolar cells of the INL. (k) Slm1 and Slm2 expression in TYW3 homozygous retina. While Slm1 levels decrease, there does not appear to be a significant change in the pattern of Slm2 expression. Scale bars are 40μm.

Because Slm2 is upregulated in the brains of *Khdrbs2* knock out mice (Traunmuller et. al, 2014), we investigated Slm2 expression in TYW3 homozygotes. In wild-types, Slm2 was expressed by retinal ganglion, amacrine, and bipolar cells (Fig. 9j). We found no detectable upregulation of Slm2 in TYW3+/+ retinas (Fig. 9k), but its expression in most Slm1-positive cells even in wild-types suggests that it may be able to compensate for loss of Slm1 in homozygotes.

## DISCUSSION

Many researchers have benefited from transgenic mice in which reporters are expressed in specific cell types that were not readily predictable based on the expression of the gene from which the transgene’s regulatory elements were derived – in other words, transgenes exhibiting what was presumed to be insertion site-dependent expression (for example, Cohen-Tannoudji et al., 1994; Huberman et al., 2009; Haverkamp et al., 2009; Trenholm et al., 2011; Kay et al., 2011b; Dhande et al., 2013; Krishnaswamy et al., 2015; Peng et al., 2017). Although identifying endogenous genes near the transgene could aid in interpreting transgene expression patterns, this has been attempted infrequently, in large part because methods for determining insertion sites have been unreliable. We were motivated to reexamine this issue for two reasons. First, initial reports suggested that the TLA method would be more reliable than its predecessors (de Vree et al., 2014; Cain-Hom et al., 2017). Second, in the course of our developmental studies, we obtained suggestive evidence that two such transgenes were inserted in close proximity to genes expressed in small neuronal subsets that the transgenes marked: HB9-GFP near *Cdh6* and Mito-P near *Fat4.* Results reported here confirm those suppositions, demonstrate linkage of the TYW3 transgene to *Khdrbs2* and, more important, provide new insights into the influence of transgenes and endogenous genes on each other.

### Endogenous genes affect expression of neighboring transgenes

Our claim for an effect of endogenous genes on transgene expression is based on the selective expression of the HB9-GFP transgene in Cdh6-positive V-ooDSGCs and the selective expression of the Mito-P transgene in Fat4-positive BC1 bipolar cells. Although we cannot entirely rule out the possibility that the correspondence is coincidental, it is highly unlikely. Of >120 retinal cell types, 3 express *Cdh6* (V-ooDSGCs, D-ooDSGCs, and starburst amacrine cells) and 2 are GFP-positive in HB9-GFP transgenic mice (V-ooDSGCs and cones). The odds of one of the GFP-positive types being Cdh6-positive by chance are 0.05 (see Methods). Likewise, 2 cell types are Fat4-positive (BC1A and BC1B) and 3 are CFP-positive in the Mito-P line (BC1A, BC1B and nGnG amacrines), with the odds of two CFP-positive types being *Fat4* positive by chance being 0.0004. This level of co-expression is therefore unlikely to be random.

Two aspects of the influence of the insertion site are noteworthy. First, as detailed above, it is incomplete, differing from patterns seen in “enhancer traps” that often faithfully report on the expression of a neighboring gene. What other factors might influence transgene expression? One possibility is that it reflects expression of the gene from which the transgene’s regulatory elements are derived, but this seems unlikely. *Hb9* is not detectably expressed in RGCs or cones (Peng et. al, 2017; Sarin et. al, 2018) and although some bipolar cells express *Thy1,* they do so at substantially lower levels than RGCs (Barnstable and Drager, 1984; Macosko et al., 2015). Another possibility is that rearrangements within the transgene or at the insertion site have generated new specificities, such as the deletion directly upstream of the Mito-P transgene and the duplication within the HB9-GFP transgene (Figures 3 and 4). Yet another, and perhaps most likely, is the one initially proposed for unexpected patterns of transgene expression, that juxtaposition of sequences within and outside of the transgene generates novel specificities (Palmiter et al., 1983)

A second point of interest is that the distances between these two transgenes and the nearest annotated genes are rather large: HB9-GFP is ~680 kb from *Cdh6* and Mito-P is ~550 kb downstream of *Fat4.* Although such long-distance interactions were once thought to be unusual, recent studies have shown that chromatin is organized into regions such as topologically associating domains or TADs, ranging from a few hundred kilobases to a few megabases, within which gene expression is coordinated by enhancers that act over the entire domain (Symmons et al., 2014; Dekker and Heard, 2015; Dixon et al., 2016). Using the dataset of Dixon et al. (2016; Wang et. al, 2017), we find that HB9-GFP and *Cdh6* are indeed within the same TAD, as are Mito-P and *Fat4* (Fig. 10). This 3-dimensional genomic compartmentalization produces secondary and tertiary structures, leading to interactions between regions that are separated by substantial linear sequence. Moreover, both transgenes are inserted in gene-poor regions, with no other annotated genes over spans of 2.9 Mb and 1.0 Mb surrounding HB9-GFP and Mito-P respectively (Figs. 3 and 4). This contrasts with an average intergenic distance of approximately 100 kb in the mouse genome (approximately 2×10^4^ genes in a 2×10^9^bp genome; see also, Mayer et al., 2005). In such regions, complex interfering signals from multiple genes or boundaries that insulate genes from each other may be minimized. Thus, while the HB9-GFP and Mito-P transgenes are hundreds of kilobases from their endogenous neighbors, gene-poor regions adjacent to the transgenes and their insertion into TADs may explain the selective expression of these transgenes by cell types also expressing their nearest neighbors.

**Figure 10.**
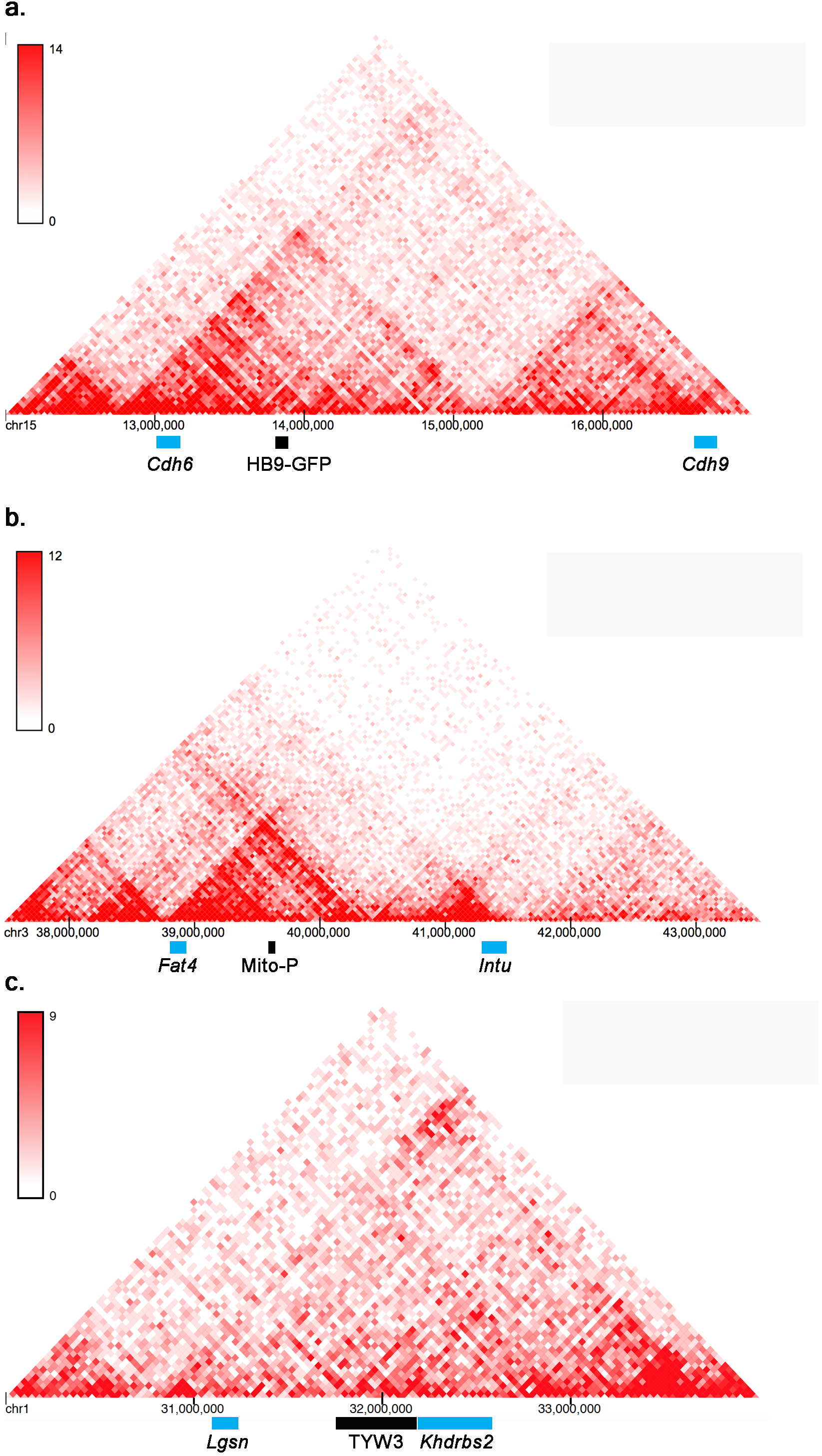
Topologically associating domains (TADs) surrounding HB9-GFP, Mito-P, and TYW3. Predicted topologically associating domains (conserved regions of 3-dimensional interaction) surrounding *Cdh6* on Chromosome 15 (a), (b) *Fat4* on Chromosome 3 (b), and *Khdrbs2* on Chromosome 1 (c), calculated using the tools in Wang et al. (2017). *Cdh6* and HB9-GFP appear in the same TAD, as do *Fat4* and Mito-P. There are no other genes within these TADs. The insertion site of TYW3 does not appear to be within a particularly interactive region of Chromosome 1. Individual bins represent 40 kb.

In contrast to *Cdh6*/HB9-GFP and *Fat4*/Mito-P, it is uncertain whether the TYW3 expression pattern is influenced by the nearby *Khdrbs2* for several reasons. *Thy1* is already expressed by all RGCs in wild-type retina and most transgenes incorporating *Thy1* regulatory elements are expressed by at least some RGCs (Feng et al., 2000; Kim et al., 2010; Misgeld et al., 2007). Likewise, most RGCs are *Khdrbs2*-positive, and YFP-positive W3-RGCs did not express *Khdrbs2* at detectably higher levels than their YFP-negative neighbors. Thus, it is not possible to disentangle the effects of *Thy1-* and *Khdrbs2-derived* regulatory sequences.

### Transgenes affect expression of neighboring endogenous genes

Many cases have been described in which insertion of a transgene mutates an endogenous gene, leading to severe defects or lethality (e.g., Soriano et al., 1987; Keller et al., 1990; Woychik and Alagramam, 1998). Indeed, some genetic screens have relied on insertional mutagenesis, using transposons and retroviral vectors as mutagens (Golling et al., 2002; Shima et al., 2016). Nonetheless, the possibility that transgenic reporters may affect endogenous genes in ways that lead to subtle defects is less often considered. It is therefore sobering that all three of the transgenes we studied affected expression of a neighboring endogenous gene. The effect, in this admittedly small sample, was distance dependent: greatest for TYW3 and *Khdrbs2,* separated by 7 kb, modest for Mito-P and *Fat4,* separated by 550 kb and small but significant for HB9-GFP and *Cdh6,* separated by 680 kb. Interference with the endogenous gene might result from interruption of endogenous regulatory elements, the deletions and rearrangements that accompany insertion, alterations in chromatin structure, or some combination.

For TYW3, the decrease in *Slm1* expression was striking. We observed no overt phenotype, consistent with the finding that even *Slm1* null mutants are viable, fertile and outwardly normal (Iijima et al., 2014 Traunmüller et al., 2014). Nonetheless, Khdrbs2 (Slm1) and its homologues, Khdrbs1 (Sam68) and Khdrbs3 (Slm2) are known to regulate alternative splicing of critical neuronal genes (Ehrmann, et. al, 2016) so alternations in its activity could affect neuronal development or function. Likewise, *Fat4* has been implicated in several developmental processes, with the null mutant being neonatally lethal (Saburi et. al, 2008). Thus, even the modest defects observed in Mito-P could be consequential under some circumstances.

### Mapping mouse transgene integration sites is feasible and useful

We have demonstrated complex reciprocal interactions between transgenes and neighboring endogenous genes. These interactions are interesting, but also potentially worrisome, since the endogenous gene affected by the transgene has a substantial chance of being expressed in the very cells that the transgene is being used to study.

We analyzed only three lines and we deliberately chose ones that exhibited interesting integration site-dependent expression patterns in retina, so it is difficult to draw strong conclusions about the frequency of these interactions. Nonetheless, there are several reasons to believe that they are more frequent than has been generally appreciated. First, we observed effects of an endogenous gene on transgene expression in two and possibly all three of the lines, and effects of the transgene on expression of the neighboring endogenous gene in all three of the lines. Second, for two of the three lines, the distance between transgene and nearest endogenous neighbor is >500 kb, dispelling the notion that insertion within a gene is prerequisite to interaction. Third, sporadic reports appearing over a long period have provided additional cases in which transgene insertions near but not within endogenous genes affect expression of the endogenous gene (e.g., Sharpe et. al, 1999; Mukai et al., 2006), or the endogenous gene influences expression of the transgene (e.g., Kothary et al., 1988, Sharpe et. al, 1999; Narboux-Nême et al., 2012).

Together, our results show that mapping of transgene insertion sites can be useful in at least three respects: First, once neighboring genes have been identified, their expression can be assayed to test the possibility that a transgenic line is in fact a hypomorph. Second, if the endogenous gene is expressed in cells marked by the transgene, it becomes a candidate effector of that cell’s development or function. Third, once the insertion site has been mapped, it becomes straightforward to devise genotyping protocols that are specific to the line and that readily distinguish heterozygotes from homozygotes. Additional potential uses include targeting new transgenes to insertion sites expected to confer desirable expression patterns on them.

To date, mapping of transgene insertion sites has not been standard practice, both because its value has been questionable and reliable methods for doing so have not been available. With improved methods now available and evidence accumulating that interactions of transgenes and endogenous genes are frequent, it may be advisable to make this a more common practice.

## AUTHOR CONTRIBUTIONS

XD, ML, MQ and IW performed experiments and analyzed data. JS conceived the project and analyzed data. ML and JS wrote the manuscript. All authors reviewed and edited the manuscript

## ACKNOWLEDGEMENTS

We thank Karthik Shekhar for statistical help; Max van Min and Judith Bergboer (Cergentis) for assistance in interpreting TLA results and permission to use material from their reports in Figures 3–5; Helen McNeil (Lunenfeld Institute) for providing the Fat4 conditional mutant and Peter Scheiffele. (Biozentrum, Basel) for antibodies to Slm1 and Slm2.

## REFERENCES

Barnstable CJ, Dräger UC. Thy-1 antigen: a ganglion cell specific marker in rodent retina. Neuroscience. 1984;11:847–55.

Bassett EA, Pontoriero GF, Feng W, Marquardt T, Fini ME, Williams T, West-Mays JA. Conditional deletion of activating protein 2alpha (AP-2alpha) in the developing retina demonstrates non-cell-autonomous roles for AP-2alpha in optic cup development. Mol Cell Biol. 2007; 27:7497–7510.

Bier E, Vaessin H, Shepherd S, Lee K, McCall K, Barbel S, Ackerman L, Carretto R, Uemura T, Grell E, Jan LY and Jan YN. Searching for pattern and mutation in the Drosophila genome with a P-lacZ vector. Genes Dev. 1989; 3:1273–87.

Brinster RL, Palmiter RD. Introduction of genes into the germ line of animals. Harvey Lect. 1984-1985;80:1–38.

Brinster RL, Chen HY, Trumbauer M, Senear AW, Warren R, Palmiter RD. Somatic expression of herpes thymidine kinase in mice following injection of a fusion gene into eggs. Cell. 1981; 27:223–31.

Burgess DL, Kohrman DC, Galt J, Plummer NW, Jones JM, Spear B, Meisler MH. Mutation of a new sodium channel gene, Scn8a, in the mouse mutant ‘motor endplate disease’. Nat Genet. 1995; 10:461–5.

Cain-Hom C, Splinter E, van Min M, Simonis M, van de Heijning M, Martinez M, Asghari V, Cox JC, Warming S. Efficient mapping of transgene integration sites and local structural changes in Cre transgenic mice using targeted locus amplification. Nucleic Acids Res. 2017; 45:e62.

Cohen-Tannoudji M, Babinet C, Wassef M (1994) Early determination of a mouse somatosensory cortex marker. Nature 368:460–463.

Dekker J, Heard E. Structural and functional diversity of Topologically Associating Domains. FEBS Lett. 2015;589:2877–84.

de Vree PJ, de Wit E, Yilmaz M, van de Heijning M, Klous P, Verstegen MJ, Wan Y, Teunissen H, Krijger PH, Geeven G, Eijk PP, Sie D, Ylstra B, Hulsman LO, van Dooren MF, van Zutven LJ, van den Ouweland A, Verbeek S, van Dijk KW, Cornelissen M, Das AT, Berkhout B, Sikkema-Raddatz B, van den Berg E, van der Vlies P, Weening D, den Dunnen JT, Matusiak M, Lamkanfi M, Ligtenberg MJ, ter Brugge P, Jonkers J, Foekens JA, Martens JW, van der Luijt R, van Amstel HK, van Min M, Splinter E, de Laat W. Targeted sequencing by proximity ligation for comprehensive variant detection and local haplotyping. Nat Biotechnol. 2014;32:1019–25.

Caroni P. Overexpression of growth-associated proteins in the neurons of adult transgenic mice. J Neurosci Methods. 1997;71:3–9.

Dhande OS, Estevez ME, Quattrochi LE, El-Danaf RN, Nguyen PL, Berson DM, Huberman AD. Genetic dissection of retinal inputs to brainstem nuclei controlling image stabilization. J Neurosci. 2013; 33:17797–813.

Dixon JR, Gorkin DU, Ren B. Chromatin Domains: The Unit of Chromosome Organization. Mol Cell. 2016; 62:668–80.

Donoghue MJ, Merlie JP, Rosenthal N and Sanes JR: A rostrocaudal gradient of transgene expression in adult skeletal muscle. Proc. Natl. Acad. Sci. 1991; 88:5847–5851.

Duan X, Krishnaswamy A, De la Huerta I, and Sanes JR. Type II cadherins guide assembly of a direction-selective retinal circuit. Cell 2014; 158: 793–807.

Dubose AJ, Lichtenstein ST, Narisu N, Bonnycastle LL, Swift AJ, Chines PS, Collins FS. Use of microarray hybrid capture and next-generation sequencing to identify the anatomy of a transgene. Nucleic Acids Res. 2013; 41:e70.

Ehrmann I, Fort P, Elliott DJ. STARs in the CNS. Biochem Soc Trans. 2016;44:1066–72.

Feng G, Mellor R, Bernstein M, Keller-Peck C, Nguyen QT, Wallace M, Nerbonne JM, Lichtman JW and Sanes JR: Imaging neuronal subset in transgenic mice expressing multiple spectral variants of GFP. Neuron 2000; 28:41–51.

Golling G, Amsterdam A, Sun Z, Antonelli M, Maldonado E, Chen W, Burgess S, Haldi M, Artzt K, Farrington S, Lin SY, Nissen RM, Hopkins N. Insertional mutagenesis in zebrafish rapidly identifies genes essential for early vertebrate development. Nat Genet. 2002;31:135–40.

Gordon JW, Ruddle FH. Integration and stable germ line transmission of genes injected into mouse pronuclei. Science. 1981; 214:1244–6.

Haverkamp S, Inta D, Monyer H, Wässle H. Expression analysis of green fluorescent protein in retinal neurons of four transgenic mouse lines. Neuroscience. 2009;160:126–39.

Hottentot QP, van Min M, Splinter E, White SJ. Targeted Locus Amplification and Next-Generation Sequencing. Methods Mol Biol. 2017;1492:185–196.

Huberman AD, Wei W, Elstrott J, Stafford BK, Feller MB, Barres BA. Genetic identification of an On-Off direction-selective retinal ganglion cell subtype reveals a layer-specific subcortical map of posterior motion. Neuron. 2009;62:327–34.

Iijima T, Iijima Y, Witte H, Scheiffele P. Neuronal cell type-specific alternative splicing is regulated by the KH domain protein SLM1. J Cell Biol. 2014;204:331–42.

Kay JN, De la Huerta I, Kim IJ, Zhang Y, Yamagata M, Chu MW, Meister M, Sanes JR. Retinal ganglion cells with distinct directional preferences differ in molecular identity, structure, and central projections. J Neurosci. 2011;31:7753–62.

Kay JN, Voinescu PE, Chu MW, and Sanes JR. Neurod6 expression defines novel retinal amacrine cell subtypes and regulates their fate. Nature Neuroscience 2011a; 14:965–972.

Kay JN, Chu MW, Sanes JR. MEGF10 and MEGF11 mediate homotypic interactions required for mosaic spacing of retinal neurons. Nature. 2012;483:465–9.

Keller SA, Liptay S, Hajra A, Meisler MH. Transgene-induced mutation of the murine steel locus. Proc Natl Acad Sci U S A. 1990;87:10019–22.

Kim IJ, Zhang Y, Meister M, Sanes JR. Laminar Restriction of Retinal Ganglion Cell Dendrites and Axons: Subtype-Specific Developmental Patterns Revealed With Transgenic Markers. Journal of Neuroscience 2010; 30:1452–62.

Kothary R, Clapoff S, Brown A, Campbell R, Peterson A, Rossant J. A transgene containing lacZ inserted into the dystonia locus is expressed in neural tube. Nature. 1988;335:435–7.

Krishnaswamy A, Yamagata M, Duan X, Hong YK, and Sanes JR. Sidekick 2 directs formation of a retinal pathway that detects differential motion. Nature 2015; 524:466–70.

Liang Z, Breman AM, Grimes BR, Rosen ED. Identifying and genotyping transgene integration loci. Transgenic Res. 2008;17:979–83.

Macosko EZ, Basu A, Satija R, Nemesh J, Shekhar K, Goldman M, Tirosh I, Bialas AR, Kamitaki N, Martersteck EM, Trombetta JJ, Weitz DA, Sanes JR, Shalek AK, Regev A, McCarroll SA. Highly Parallel Genome-wide Expression Profiling of Individual Cells Using Nanoliter Droplets. Cell. 2015; 161:1202–1214.

Mayer R, Brero A, von Hase J, Schroeder T, Cremer T, Dietzel S. Common themes and cell type specific variations of higher order chromatin arrangements in the mouse. BMC Cell Biol. 2005;6:44.

Misgeld T, Kerschensteiner M, Bareyre FM, Burgess RW, Lichtman JW. Imaging axonal transport of mitochondria in vivo. Nat Methods 4: 559–561, 2007.

Mukai HY, Motohashi H, Ohneda O, Suzuki N, Nagano M, Yamamoto M. Transgene insertion in proximity to the c-myb gene disrupts erythroid-megakaryocytic lineage bifurcation Mol Cell Biol. 2006;26:7953–65.

Narboux-Nême N, Goïame R, Mattéi MG, Cohen-Tannoudji M, Wassef M. Integration of H-2Z1, a somatosensory cortex-expressed transgene, interferes with the expression of the Satb1 and Tbc1d5 flanking genes and affects the differentiation of a subset of cortical interneurons. J Neurosci. 2012;32:7287–300.

Palmiter RD, Brinster RL. Germ-line transformation of mice. Annu Rev Genet. 1986;20:465–99.

Palmiter RD, Norstedt G, Gelinas RE, Hammer RE, Brinster RL. Metallothionein-human GH fusion genes stimulate growth of mice. Science. 1983;222:809–14.

Peng YR, Tran NM, Krishnaswamy A, Kostadinov D, Martersteck EM, Sanes JR. Satb1 Regulates Contactin 5 to Pattern Dendrites of a Mammalian Retinal Ganglion Cell. Neuron. 2017;95:869–883.

Raman P, Grachtchouk V, Lyons RH Jr, Koenig RJ. Identification of the Genomic Insertion Site of the Thyroid Peroxidase Promoter-Cre Recombinase Transgene Using a Novel, Efficient, Next-Generation DNA Sequencing Method. Thyroid. 2015;25:1162–6.

Rao MV, Donoghue MJ, Merlie JP and Sanes JR: Distinct regulatory elements control muscle-specific, fiber type-selective, and axially-graded expression of a myosin light chain gene in transgenic mice. Mol. Cell. Biol. 1996; 16:3909–3922.

Ray TA, Roy S, Kozlowski C, Wang J, Cafaro J, Hulbert SW, Wright CV, Field GD, Kay JN. Formation of retinal direction-selective circuitry initiated by starburst amacrine cell homotypic contact. Elife. 2018;7. pii: e34241.

Rodriguez AR, de Sevilla Müller LP, Brecha NC. The RNA binding protein RBPMS is a selective marker of ganglion cells in the mammalian retina. J Comp Neurol. 2014;522:1411–43.

Saburi S, Hester I, Fischer E, Pontoglio M, Eremina V, Gessler M, Quaggin SE, Harrison R, Mount R, McNeill H. Loss of Fat4 disrupts PCP signaling and oriented cell division and leads to cystic kidney disease. Nat Genet. 2008;40:1010–5.

Sarin S, Zuniga-Sanchez E, Kurmangaliyev YZ, Cousins H, Patel M, Hernandez J, Zhang KX, Samuel MA, Morey M, Sanes JR, Zipursky SL. Role for Wnt Signaling in Retinal Neuropil Development: Analysis via RNA-Seq and In Vivo Somatic CRISPR Mutagenesis. Neuron. 2018;98:109–126.

Sethuramanujam S, Yao X, deRosenroll G, Briggman KL, Field GD, Awatramani GB. “Silent” NMDA Synapses Enhance Motion Sensitivity in a Mature Retinal Circuit. Neuron. 2017;96:1099–1111.

Schubert T, Kerschensteiner D, Eggers ED, Misgeld T, Kerschensteiner M, Lichtman JW, Lukasiewicz PD, Wong RO. Development of presynaptic inhibition onto retinal bipolar cell axon terminals is subclass-specific. J Neurophysiol. 2008;100:304–16.

Sha H, Xu J, Tang J, Ding J, Gong J, Ge X, Kong D, Gao X. Disruption of a novel regulatory locus results in decreased Bdnf expression, obesity, and type 2 diabetes in mice. Physiol Genomics. 2007; 31:252–63.

Sharpe J, Lettice L, Hecksher-Sorensen J, Fox M, Hill R, Krumlauf R. Identification of sonic hedgehog as a candidate gene responsible for the polydactylous mouse mutant Sasquatch. Curr Biol. 1999;9:97–100.

Shekhar K, Lapan SW, Whitney IE, Tran NM, Macosko EZ, Kowalczyk K, Adiconis X, Levin JZ, Nemesh J, Goldman M, McCarroll SA, Cepko CL, Regev A, and Sanes JR. Comprehensive classification of retinal bipolar neurons by single-cell transcriptomics. Cell, 2016; 166:1308–1323.

Shima Y, Sugino K, Hempel CM, Shima M, Taneja P, Bullis JB, Mehta S, Lois C, Nelson SB. A Mammalian enhancer trap resource for discovering and manipulating neuronal cell types. Elife. 2016;5:e13503.

Soriano P, Gridley T, Jaenisch R. Retroviruses and insertional mutagenesis in mice: proviral integration at the Mov 34 locus leads to early embryonic death. Genes Dev. 1987;1:366–75.

Srivastava A, Philip VM, Greenstein I, Rowe LB, Barter M, Lutz C, Reinholdt LG. Discovery of transgene insertion sites by high throughput sequencing of mate pair libraries. BMC Genomics. 2014;15:367.

Swanson LW, Simmons DM, Arriza J, Hammer R, Brinster R, Rosenfeld MG, Evans RM. Novel developmental specificity in the nervous system of transgenic animals expressing growth hormone fusion genes. Nature. 1985;317:363–6.

Suzuki O, Hata T, Takekawa N, Koura M, Takano K, Yamamoto Y, Noguchi Y, Uchio-Yamada K, Matsuda J. Transgene insertion pattern analysis using genomic walking in a transgenic mouse line. Exp Anim. 2006;55:65–9;

Symmons O, Uslu VV, Tsujimura T, Ruf S, Nassari S, Schwarzer W, Ettwiller L, Spitz F. Functional and topological characteristics of mammalian regulatory domains. Genome Res. 2014;24:390–400.

Traunmüller L, Bornmann C, Scheiffele P. Alternative splicing coupled nonsense-mediated decay generates neuronal cell type-specific expression of SLM proteins. J Neurosci. 2014;34:16755–61.

Trenholm S, Johnson K, Li X, Smith RG, Awatramani GB. Parallel mechanisms encode direction in the retina. Neuron. 2011;71:683–94.

Trenholm S, Johnson K, Li X, Smith RG, Awatramani GB. Erratum to Parallel mechanisms encode direction in the retina. Neuron. 2013;77:204–8.

Wang Y, Zhang B, Zhang L, An L, Xu J, Li D, Choudhary MNK, Li Y, Hu M, Hardison R, Wang T, Yue F. The 3D Genome Browser: a web-based browser for visualizing 3D genome organization and long-range chromatin interactions. https://www.biorxiv.org/content/early/2017/02/27/112268/

Weis J, Fine SM, David C, Savarirayan S and Sanes JR: Integration site-dependent expression of a transgene reveals specialized features of cells associated with neuromuscular junctions. J. Cell Biol. 1991; 113:1385–1397.

Wichterle H, Lieberam I, Porter JA, Jessell TM. Directed differentiation of embryonic stem cells into motor neurons. Cell. 2002;110:385–97.

Williams A, Harker N, Ktistaki E, Veiga-Fernandes H, Roderick K, Tolaini M, Norton T, Williams K, Kioussis D. Position effect variegation and imprinting of transgenes in lymphocytes Nucleic Acids Res. 2008; 36:2320–9.

Woychik RP, Alagramam K. Insertional mutagenesis in transgenic mice generated by the pronuclear microinjection procedure. Int J Dev Biol. 1998; 42:1009–17.

Zhang, Y, Kim, I-J, Sanes, JR, and Meister, M, The Most Numerous Ganglion Cell Type of the Mouse Retina is a Selective Feature Detector. Proc Natl Acad Sci U S A. 2012; 109:E2391–2398.

